# The role of chromatin accessibility in cis-regulatory evolution

**DOI:** 10.1101/319046

**Authors:** Pei-Chen Peng, Pierre Khoueiry, Charles Girardot, James P. Reddington, David A. Garfield, Eileen E.M. Furlong, Saurabh Sinha

## Abstract

Transcription factor (TF) binding is determined by sequence as well as chromatin accessibility. While the role of accessibility in shaping TF-binding landscapes is well recorded, its role in evolutionary divergence of TF binding, which in turn can alter cis-regulatory activities, is not well understood. In this work, we studied the evolution of genome-wide binding landscapes of five major transcription factors (TFs) in the core network of mesoderm specification, between *D. melanogaster* and *D. virilis*, and examined its relationship to accessibility and sequence-level changes. We generated chromatin accessibility data from three important stages of embryogenesis in both *D. melanogaster* and *D. virilis*, and recorded conservation and divergence patterns. We then used multi-variable models to correlate accessibility and sequence changes to TF binding divergence. We found that accessibility changes can in some cases, e.g., for the master regulator Twist and for earlier developmental stages, more accurately predict binding change than is possible using TF binding motif changes between orthologous enhancers. Accessibility changes also explain a significant portion of the co-divergence of TF pairs. We noted that accessibility and motif changes offer complementary views of the evolution of TF binding, and developed a combined model that captures the evolutionary data much more accurately than either view alone. Finally, we trained machine learning models to predict enhancer activity from TF binding, and used these functional models to argue that motif and accessibility-based predictors of TF binding change can substitute for experimentally measured binding change, for the purpose of predicting evolutionary changes in enhancer activity.

## INTRODUCTION

Cis-regulatory evolution plays an important role in phenotypic diversity, including morphological (Prud’Homme, et al. 2006), physiological (Siepel and Arbiza 2014) and behavioral (Saul, et al. 2017; Wray 2007) evolution. Given its importance, many studies have examined cross-species changes in various aspects of gene regulation, including expression (Paris, et al. 2013), enhancer activity (Khoueiry, et al. 2017), transcription factor (TF)-DNA binding (Bradley, et al. 2010; Carvunis, et al. 2015; Stefflova, et al. 2013; Wong, et al. 2015), TF binding motifs (Bradley, et al. 2010; Cheng, et al. 2014; He, et al. 2011; Moses, et al. 2006; Naval-Sánchez, et al. 2015), DNA accessibility (Alexandre, et al. 2017) and chromatin states (Lesch, et al. 2016). Changes have been observed to differing extents in these measurable aspects of gene regulation, leading to some emerging principles underlying their conservation and divergence (Cheng, et al. 2014). At the same time, it is challenging to systematically integrate these diverse qualitative observations about *cis*-regulatory evolution at different regulatory levels given the different phyla, biological systems, and technologies.

A number of studies have examined the evolution of DNA cis-regulatory sequences (Garfield, et al. 2012; Wittkopp and Kalay 2012). Some have noted surprisingly high levels of sequence change (Doniger and Fay 2007; Emberly, et al. 2003; Yokoyama, et al. 2014), but regulatory function and gene expression are often conserved despite sequence-level changes (Arnold, et al. 2014; Duque, et al. 2013; Khoueiry, et al. 2017; Ludwig, et al. 2000; Yang, et al. 2015), revealing considerable flexibility in sequence encoding the same function (Hare, et al. 2008; Kazemian, et al. 2014). Further investigations asked if the observed functional buffering against sequence divergence happens at the level of TF-DNA binding, which is the principle molecular event mediating sequence-expression relationships. ChIP-chip or ChIP-seq assays of the same TFs were performed in multiple species (Bradley, et al. 2010; Carvunis, et al. 2015; Khoueiry, et al. 2017; Stefflova, et al. 2013) and while genome-wide TF binding landscapes were noted to be conserved overall, many large qualitative as well as quantitative differences in binding were also reported (Bradley, et al. 2010). The evolution of binding landscapes thus emerged as an intriguing aspect of molecular evolution, and researchers sought to identify its main determinants.

Loss and gain of TF ChIP peaks are correlated with changes in the presence of the TF’s DNA binding motif (Bradley, et al. 2010; Cheng, et al. 2014; He, et al. 2011; Moses, et al. 2006; Naval-Sánchez, et al. 2015), but this relationship, though significant in its extent, was far from a satisfactory explanation for TF binding differences. For instance, many peaks are lost though the motif is conserved, and conversely, peaks are often conserved despite motif loss. Some studies noted the influence of co-binding TFs at or near the peak (Stefflova, et al. 2013), suggesting roles for co-operative (Duque, et al. 2013) and ‘TF collective’ modes of occupancy (Khoueiry, et al. 2017).

On the other hand, Bradley et al. (2010)observed a correlation between evolutionary changes of occupancy among multiple TFs and interpreted this as evidence for TF-independent influences such as differences in local chromatin accessibility. Indeed, DNA accessibility is known to be a major correlate of TF-DNA binding in individual species (Cheng, et al. 2013; Li, et al. 2011), and may therefore underlie evolutionary changes in TF binding. For instance, Paris et al. (2013) noted that binding divergence is correlated with changes in binding sites for the pioneer factor Zelda, which indirectly implicates accessibility changes. Genome-wide accessibility landscapes are generally evolutionarily conserved (Connelly, et al. 2014; Vierstra, et al. 2014), but accessibility changes between orthologous genomic elements are also observed and raise the question: how often do they underlie evolutionary changes in TF binding? Surprisingly, there is no direct analysis of this question. In related work, Connelly et al. (2014) reported evidence that much of accessibility divergence (between two yeast species) may be inconsequential for gene expression. Alexandre et al. (2017) made similar observations for different ecotypes of A. thaliana, but also noted that loci with high sequence variation and accessibility changes were significantly linked to expression changes. However, the extent to which accessibility changes are predictive of TF binding changes between species remains unknown. Is this relationship comparable in extent to the documented relationship between motif change and TF binding divergence? Do changes in accessibility and motif presence carry complementary information related to observed changes in TF ChIP peaks? How often is accessibility conserved, yet a TF’s occupancy diverged due to motif turnover, and how commonly do changes in accessibility result in loss or gain of TF ChIP peaks despite conservation of motif presence? These are not mutually exclusive possibilities and teasing apart their relative contributions and potential causal influence requires a formal, quantitative analysis. Insights emerging from such analyses may also fuel discussions of cause-versus-effect in the relationship between TF binding and accessibility (Guertin and Lis 2013; Li, et al. 2011). In addition to advancing our basic understanding of *cis*-regulatory evolution, answering these questions may also allow us to predict changes in TF binding using computational models that incorporate data on sequence and accessibility changes, bypassing the need for expensive ChIP profiling of TFs across species and individuals.

Any investigation of these aspects of *cis*-regulatory evolution must also consider promiscuous occupancy of TFs (Li, et al. 2008) and that a large number of ChIP peaks may have no functional impact on gene expression (Cusanovich, et al. 2014), or be functionally redundant. Evolutionary comparisons have strongly suggested that expression changes are poorly explained by TF binding changes (Paris, et al. 2013), underscoring the need to examine evolutionary questions about TF binding in a functional context. It is difficult to generally predict whether a TF ChIP peak is functional, but there are a few well-characterized regulatory systems where detailed prior knowledge of the regulatory network permits such an exercise. One of these systems is the mesoderm specification network in *D. melanogaster*, where extensive prior work has established the role of a small set of TFs in determining spatio-temporal expression patterns of a large number of genes (Azpiazu and Frasch 1993; Baylies and Bate 1996; Jakobsen, et al. 2007; Jin, et al. 2013; Khoueiry, et al. 2017; Sandmann, et al. 2007; Sandmann, et al. 2006; Yin and Frasch 1998; Yin, et al. 1997; Zaffran, et al. 2001; Zinzen, et al. 2009). This has previously led to the cataloging of thousands of putative enhancers responsible for such patterning (Khoueiry, et al. 2017; Zinzen, et al. 2009), with hundreds of them being experimentally validated through reporter assays in transgenic embryos. The mesoderm network with its richness of prior knowledge and *cis*- regulatory data sets thus provides a uniquely suited system to investigate cross-species evolution of TF binding and its determinants (Khoueiry, et al. 2017).

In this work, we studied the evolution of genome-wide binding landscapes of five essential TFs in the mesoderm specification network, between two drosophilids *D. melanogaster* and *D. virilis*, species separated by 40 million years (Paris, et al. 2013) (1.4 substitutions per neutral site (Stark, et al. 2007)). We generated *DNase I* hypersensitive sites (DHS) data to measure chromatin accessibility at three different temporal stages during early embryonic development in both *D. melanogaster* and *D. virilis*, and recorded conservation and divergence patterns. We built predictive models that use either motif change or accessibility change to predict stage-specific binding divergence of all five TFs, using our previously reported inter-species ChIP data (Khoueiry, et al. 2017; Zinzen, et al. 2009). Using these models and focusing on a large set of previously characterized mesoderm enhancers (Khoueiry, et al. 2017; Zinzen, et al. 2009) to increase functional relevance, we found that accessibility and TF binding motif changes have comparable predictive relationship with changes in TF binding. We also noted that they bear complementary information and showed that a model using both accessibility and motif information can predict TF binding divergence with significantly greater accuracy than models using either type of information alone. All of these analyses were conducted separately for each TF. In a final analysis, we used machine learning models to examine changes in TF binding of multiple factors simultaneously, in terms of their combinatorial effects on gene expression. We found that motif and accessibility based predictors of TF binding change can substitute for experimentally measured binding change, for the purpose of predicting divergence in gene expression.

## RESULTS

### Evolutionary changes in TF-DNA binding and DNA accessibility in the context of a well-characterized regulatory network

To understand how evolutionary changes of sequence and accessibility affect TF binding and enhancer activities, we focused our study on an extensively studied regulatory network where prior knowledge of essential regulators and functional enhancers can effectively guide us to functional TF-DNA binding events. We analyzed TF occupancy data for five TFs that form the core of a regulatory network essential for mesoderm development in *Drosophila* (Wilczynski and Furlong 2010): Twist (Twi), Myocyte enhancer factor-2 (Mef2), Tinman (Tin), Bagpipe (Bap) and Biniou (Bin) (Figure 1A). We obtained genome-wide TF-DNA binding information on these five TFs across five stages of embryonic development (henceforth, ‘time points’ or ‘TP’s), in the form of ChIP-chip and ChIP-seq assays in *D. melanogaster* (Zinzen, et al. 2009) and *D. virilis (Khoueiry, et al. 2017)* respectively. A total of 14 TF-time point pairs (Figure 1B), henceforth called ‘TF:TP conditions’ or simply ‘conditions’, were represented in the data, originally reported in our previous work (Khoueiry, et al. 2017). To supplement these data, we also collected stage-matched DNase-Seq libraries from both *D. virilis* and *D. melanogaster* in three of the five time points, i.e. TP1, TP3, and TP5 (Figure 1B; see Methods). Over 2,500 pairs of putative orthologous enhancers involved in mesoderm specification were identified in our previous study (Khoueiry, et al. 2017), based on presence of ChIP peaks for the core TFs, and served as the targets of our computational analyses in this work. Each enhancer (in either species) was assigned a ChIP score for each TF:TP combination, combining ChIP peaks located within the same *cis*-regulatory element (see Methods).

**Figure 1.**
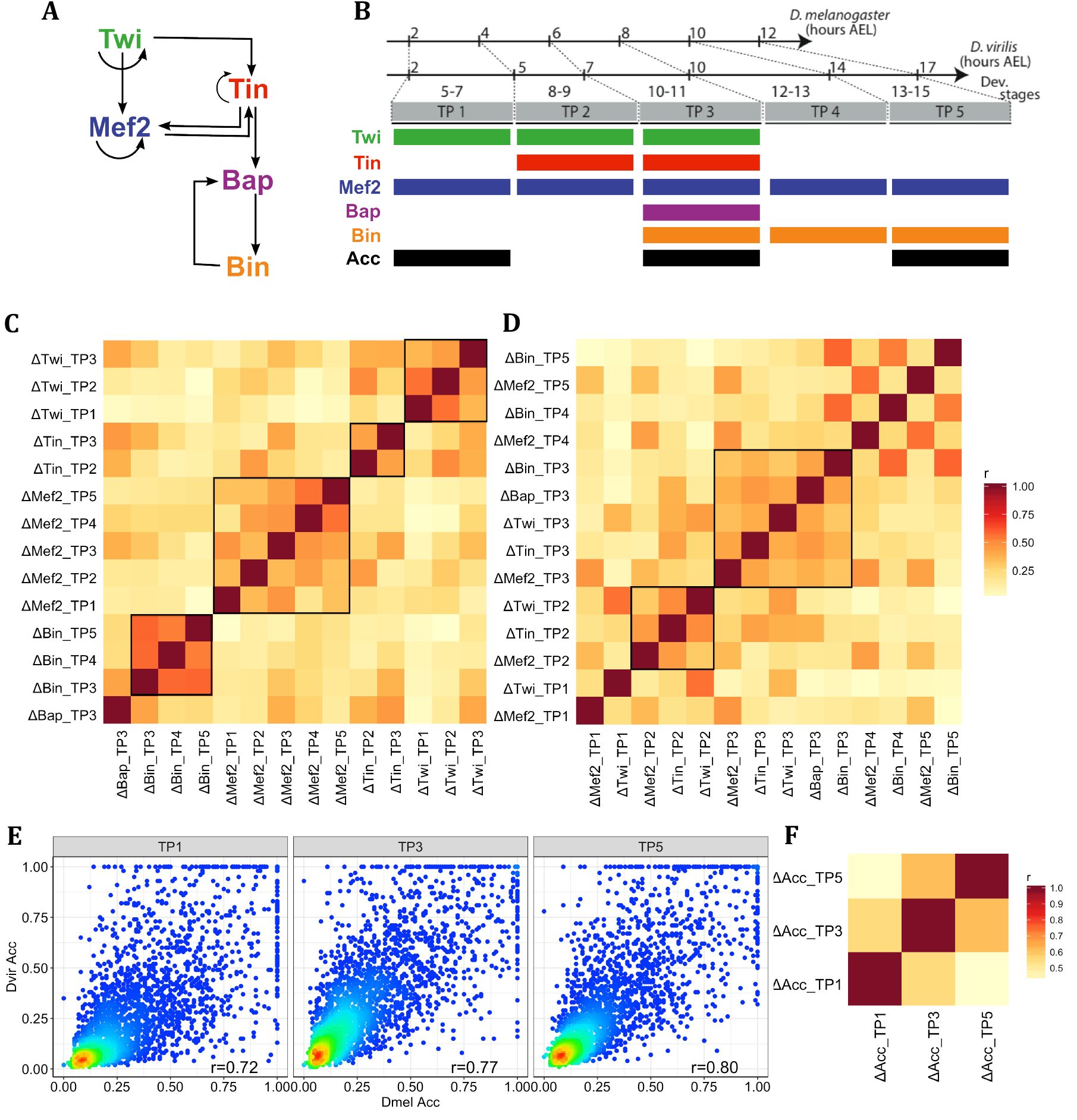
Examining evolutionary changes in TF binding and accessibility across developmental time points. (A) Regulatory network of five key TFs in mesoderm specification, source: [Khoueiry et al., eLife 2017]. (B) Data from *D. melanogaster* and *D. virilis* TF ChIP and DNase I hypersensitivity assays were collected; *D. virilis* DHS data were generated for this study. Colored boxes indicate time points (TP1-5) for which each type of genomic profile is available. Orthologous developmental stages between species were mapped according to hours of development in each species, after egg laying (AEL). (C,D) Pairwise Pearson correlations of interspecies ChIP changes, sorted by TF (C) or by time points (D) (E) Normalized accessibility scores of orthologous enhancers for three time points (TP1,3,5). Colors indicate point density, with warmer colors denoting greater density. Pearson correlations between *D. melanogaster* TF ChIP and *D. virilis* TF ChIP are also shown. (F) Pairwise Pearson correlations of interspecies accessibility changes. Data and analysis shown in (C-E) pertain to over 2,500 pairs of putative orthologous enhancers involved in mesoderm specification as defined in text.

We calculated evolutionary changes in TF occupancy as the difference of normalized quantitative ChIP scores between orthologous enhancers, for each TF at each TP (‘ΔTF:TP’). We noted extensive correlations among different ΔTF:TP measures (Figure 1C), i.e., evolutionary changes of TF-DNA binding profiles are correlated. This is especially true of binding profiles of the same TF at different time points, i.e., if a TF loses binding at a location, it tends also to lose binding at the same location at a different developmental stage. For example, Pearson correlation coefficient (PCC) of ΔBin:TP3 (changes in Bin binding at TP3) and ΔBin:TP4 is 0.58 (p-value=2.30E-247), and that between ΔMef2:TP4 and ΔMef2:TP5 is 0.56 (p-value=3.66E-227). The natural explanation for this observation is that loss or gain of the TF’s motif plays a significant role in evolutionary changes of TF binding. More interestingly, changes in DNA binding of different TFs at the same time point also show correlations (Figure 1D), e.g., ΔTin:TP2 and ΔTwi:TP2 have a Pearson correlation of 0.50 (p-value=3.69E-174). Since the five TFs have different binding preferences (motifs, see Supplementary Figure S1), these correlations most likely arise due to co-binding of specific pairs of TFs – a possibility that we examined in (Khoueiry, et al. 2017), or from changes in accessibility, which is a common contributing factor to DNA binding profiles of different TFs (Connelly, et al. 2014; Vierstra, et al. 2014).

We also compared the normalized DHS accessibility scores (see Methods) of the same set of ~2,500 orthologous enhancers mentioned above, at each of the three time-points (Figure 1E). Most of the accessibility scores are conserved between species, while some enhancer pairs exhibit substantial change. For instance, at TP1, of all the enhancer pairs whose accessibility score is above 0.3 (median score, on a scale of 0 to 1) in at least one of the two species, ~7% have their orthologous accessibility score below 0.1. We also compared evolutionary changes in accessibility between orthologous enhancers at different developmental stages and found, as expected, that the temporally proximal time points, e.g., TP3 and TP5, or TP1 and TP3, have more correlated evolutionary changes than more separated time points, i.e., TP1 and TP5 (Figure 1F). In short, we collated data on TF binding and DNA accessibility at the same stages of embryogenesis in two species, and confirmed previous reports of evolutionary flux in these important measures of the *cis*-regulatory landscape, setting the stage for a closer examination of their mutual relationship.

### Relationship between changes in chromatin accessibility and TF binding at orthologous developmental enhancers

We sought to systematically and quantitatively dissect how evolutionary changes in TF binding are related to changes in accessibility. Given the observations above, that the occupancy of different TFs from the same time point tend to change concordantly, it was natural to ask: “how frequently do changes in TF binding between species coincide with changes in DNA accessibility?” We collected orthologous enhancer pairs that are accessible in at least one of the two species (normalized accessibility score > 0.3), and examined the relationship between change of accessibility score (‘ΔAcc’) and change of TF occupancy. As shown in Figure 2A, for Twi binding at TP1, enhancers with conserved binding (points closer to diagonal) typically have conserved accessibility (warmer colors), while enhancers with changes of TF binding (off-diagonal points) tend to exhibit changes in their accessibility score (cooler colors) (Pearson correlation between ΔAcc and ΔChIP is r=0.12, p-value 1.44E-5). Other TF:TP combinations showed the same trend (Supplementary Figure S2).

**Figure 2.**
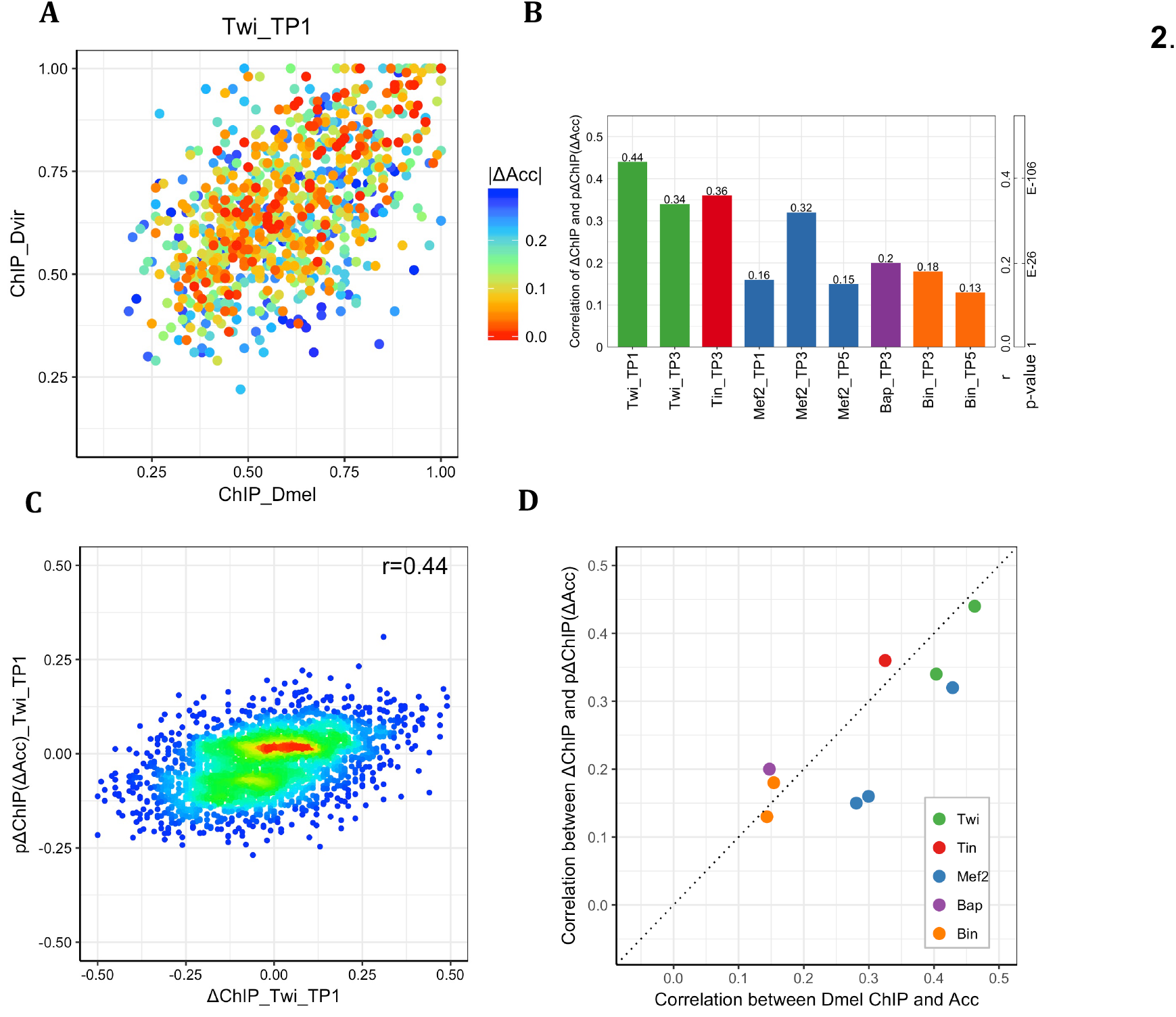
Accessibility changes alone are modest predictors of TF occupancy changes between species. (A) Scatter plot of *D. melanogaster* ChIP scores versus *D. virilis* ChIP scores for Twi at TP1. Points represent orthologous enhancers that are accessible in at least one species. Colors indicate change of accessibility score. (B) Correlation coefficient between measured ΔChIP and ΔChIP predicted based on ΔAcc, denoted as ‘pΔChIP(ΔAcc)’. P-values of Pearson correlation coefficient (r) with sample size of 2754 are also shown. (C) Scatter plot of ΔChIP versus pΔChIP for Twi at TP1. Warmer colors indicate greater point density. (D) Correlation between ChIP and accessibility in *D. melanogaster* (x-axis) is compared to correlation between interspecies ΔChIP and pΔChIP(ΔAcc).

While the above observations were statistically significant, the strength of relationship between accessibility and TF binding changes revealed by them seemed modest. In part, this may be because the quantified change in TF binding depends not only on the change of accessibility (ΔAcc) but also on the actual accessibility in either species. Thus, to make the above analysis more systematic, for each of a set of 2,754 orthologous putative enhancer pairs and for each TF:TP condition, we trained a Support Vector Regression (SVR) model to predict interspecies differences in ChIP scores (ΔChIP) using the *D. melanogaster* accessibility score and ΔAcc as features. Goodness of fit was measured by Pearson correlation coefficient between measured and model-predicted ΔChIP values, using 5-fold cross-validation. We found that changes in accessibility are modestly predictive of changes in TF binding between species, with correlation coefficients varying substantially across the 9 data sets, averaging about 0.25 (Figure 2B). To provide an intuitive calibration of this value, we note that it was computed over 2,754 samples and has a p-value of 1.64E-40. As an alternative evaluation of the predictions, we asked how well the model-predicted ΔChIP values classify the enhancer pairs with the greatest increase in TF binding (measured ΔChIP in top 10 percentile among all 2,754 orthologous pairs) versus those with the greatest decrease in binding (ΔChIP in bottom 10 percentile). We noted an AUROC of 0.78 or greater on such balanced data sets for four of the 9 TF:TP pairs (Supplementary Figure S3). This result shows that accessibility comparison between orthologs can discern cases of most extreme increase versus decrease of TF binding.

Among the best examples was Twi:TP1, where correlation between measured and predicted ΔChIP on the full set of 2,754 orthologous pairs is 0.44 (p-value 8.94E-131), i.e., about 20% of the variance (r^2^ = 0.19) of ΔChIP is explained by accessibility changes for this condition (Figure 2C). What mechanisms might underlie this relationship? An intriguing but untested possibility is that Twi is the major factor dictating open chromatin i.e., perhaps having a pioneering role at these sites, in keeping with its role as a ‘master regulator’ being sufficient to convert cells to a mesodermal fate (Baylies and Bate 1996). Alternatively, there may be unmeasured changes in an additional factor required to open chromatin and facilitate Twi binding to these sites. Zelda is a very good candidate, as it is required for Twi binding to some early developmental enhancers (Yáñez-Cuna, et al. 2012) and is thought to play a pioneer role in early *Drosophila* embryogenesis (Harrison, et al. 2011; Liang, et al. 2008; Schulz, et al. 2015). Such mechanistic speculations notwithstanding, the above results – that even in the best example only 20% of variance is explained – emphasize the potential existence of influences other than accessibility, and that operate without major effects on accessibility, on TF binding change.

A natural comparison point for the above correlations is the extent to which accessibility score in a single species can predict TF binding in that species in the same time point, across the same set of enhancers as above. It was not *a priori* clear what the result of this comparison might be. It was possible that accessibility changes are less prominently associated with evolutionary changes of TF binding, compared to the extent to which accessibility is associated with TF binding in a single species (Connelly, et al. 2014; Vierstra, et al. 2014), for instance if most binding changes arise from turnover of motif hits. On the other hand, the single species correlation values between accessibility and TF binding might not be as high here as reported in some previous studies (Cheng, et al. 2013; Li, et al. 2011), since our analysis is restricted to putative enhancers, which as a class have high accessibility levels. Our single-species correlation analysis will not reflect the strong genome-wide trend of ChIP peaks coinciding with accessible regions. With these two considerations in mind, it was thus instructive to find that correlations between ChIP score and accessibility score in *D. melanogaster* (Figure 2D) were similar to those between evolutionary changes in these scores, i.e., ΔChIP and predicted ΔChIP based on accessibility, for every TF:TP condition.

### Accessibility changes partly explain co-divergence of binding by pairs of transcription factors

We noted above (Figure 1D) that changes in binding for some TF pairs in the same time point, are strongly correlated. We asked if these co-divergence patterns can be explained by changes in accessibility, since accessibility can be simplistically thought of as setting up a ‘landscape’ for binding, on which different TFs act differently to set up their own binding profiles. Evolutionary changes in accessibility can therefore be expected to impact binding of multiple TFs in similar ways. To test this possibility, we computed a statistic similar to the partial correlation of ΔChIP between each pair of TFs, given accessibility data. For each pair of TFs, we first computed the residuals of accessibility-based predictors of ΔChIP for either TF, and then calculated the correlation coefficient between these residuals. This approach removes the effect of accessibility changes in assessing the correlation of ΔChIP between TF1 and TF2. We found that for the majority of data set pairs (10 out of 16) where ΔChIP scores of two TFs at the same time point are strongly correlated (PCC > 0.2), correlations are lower (a difference of at least 0.04) after excluding the influence of accessibility (Supplementary Table S1), though the (partial) correlations remain strong even after accounting for ΔAcc. For Twi and Tin at TP2, for example, the correlation of ΔChIP scores drops from 0.5 to 0.45 upon ‘removing’ accessibility; a similar reduction is observed for the same pair of TFs at TP3, where the correlation drops from 0.39 to 0.32. Another example is that of Mef2 and Tin at TP2, where the correlation reduced from 0.45 to 0.37 upon accounting for accessibility changes. Indeed, previous work reported the potential role of Tin-Twi and Tin-Mef2 co-binding in the evolution of binding sites for these TFs (Khoueiry, et al. 2017). Our results reveal that changes in accessibility do explain part of the co-divergence of DNA binding exhibited by pairs of TFs, but other causes of co-divergence (Duque and Sinha 2015; Khoueiry, et al. 2017), e.g., cooperative occupancy, functional change of an enhancer and the resulting shared changes of selective pressure, also exist.

### Changes in accessibility and sequence predict TF binding changes to similar extents

The results above quantified the extent to which change of accessibility (ΔAcc) predicts changes in TF-DNA binding (ΔChIP) between orthologous enhancers. We next determined how strongly changes in sequence, in terms of binding motif presence, predict ΔChIP, with the ultimate goal of comparing the relative contributions of changes in accessibility and in sequence to divergence of TF binding. To approach this goal, it is important to have a means of quantifying a TF’s motif presence in a given sequence accurately enough to allow quantitative assessment of motif change between orthologous enhancers. We used our previously published STAP (Sequence To Affinity Prediction) model (He, et al. 2009) for this purpose. STAP is a thermodynamics-based model that integrates one or more strong as well as weak binding sites, using a given motif, to predict net TF occupancy within a DNA segment. The STAP score is a more realistic estimation of motif presence in a window, compared to using the strength of the best motif match or counting the number of matches above a threshold. Importantly, it is not a confidence score of a single binding site (e.g., CENTIPEDE (Pique-Regi, et al. 2011)) and is thus better suited to assess net sequence change in developmental enhancers, which often exhibit homotypic site clustering (Ezer, et al. 2014; Lifanov, et al. 2003) and suboptimal sites (Crocker, et al. 2015; Farley, et al. 2015).

For each orthologous enhancer pair, we calculated STAP scores of either ortholog using a TF’s motif, and thus obtained a ‘ΔSTAP’ score quantifying the evolutionary change in motif presence for that TF. Next, we used the *D. melanogaster* STAP score and the ΔSTAP score together to predict ΔChIP for each orthologous enhancer pair, using a Support Vector Regression (SVR) algorithm, similar to what was done for accessibility scores in the previous section. This was repeated for each TF:TP condition. We found that the predicted and measured ΔChIP are modestly correlated, with average correlation coefficients in the 14 TF:TP conditions being ~0.3 (Figure 3A, each reported correlation is an average across 5-fold cross validation). It was notable that most conditions exhibited similar correlations, with 9 of the 14 yielding values between 0.27 and 0.33, and the highest correlation (0.38) seen for the Bin-TP3 and Bin-TP5 conditions. By way of calibration, we similarly computed correlations between STAP and ChIP scores in each species separately, across the same set of enhancers as above. We noted that correlation coefficients are ~0.68 for *D. melanogaster* and 0.61 for *D. virilis* (Supplementary Figure S4), on average across the 14 conditions. This assured us that STAP provides an accurate estimate of motif content, which is strongly predictive of TF occupancy, and does so in both species. However, it also highlights the poorer predictability of evolutionary changes in binding from change in sequence compared to the ability to predict binding from sequence in a single species.

**Figure 3.**
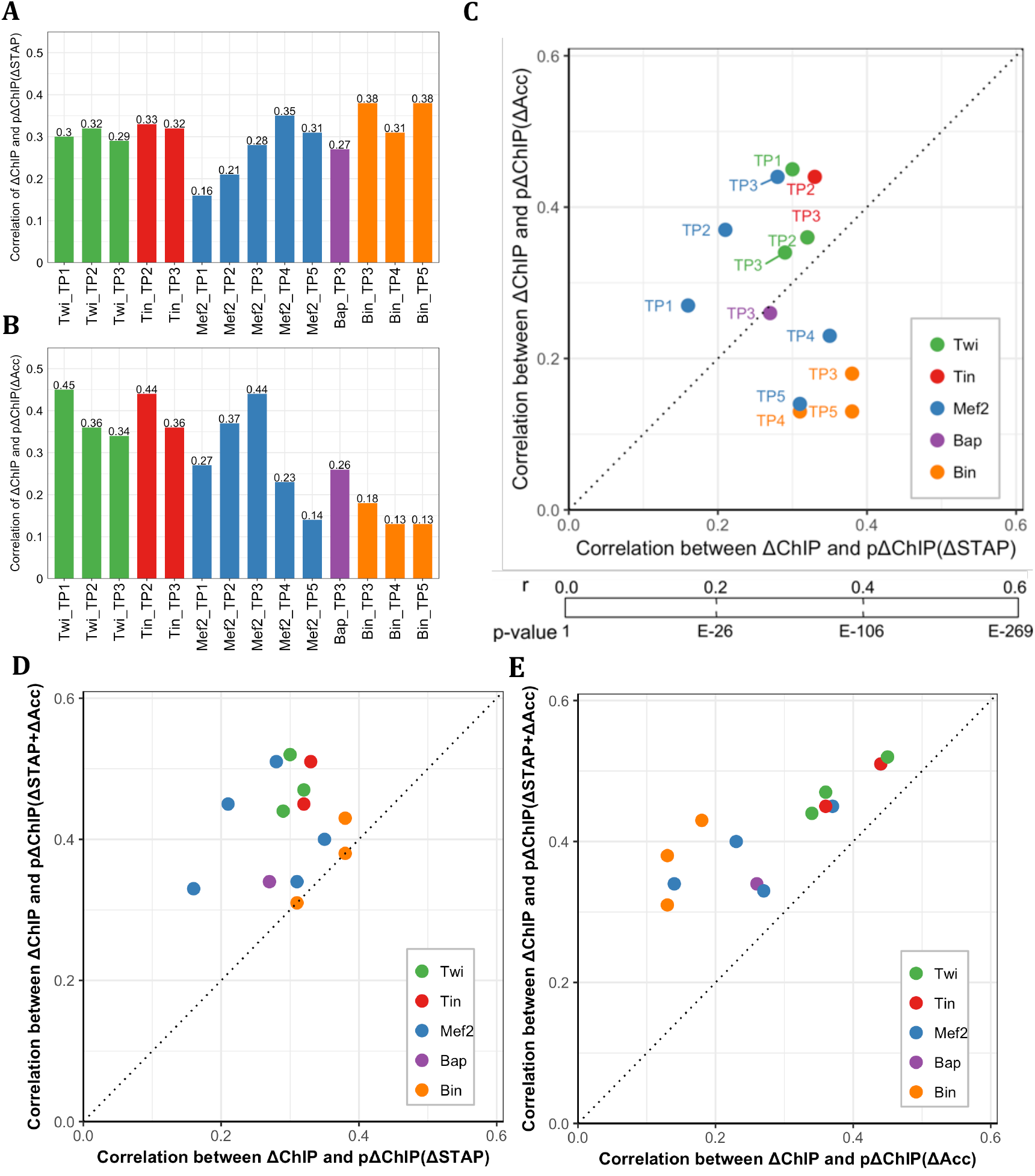
Changes in motif presence and accessibility are both predictive of TF occupancy change. (A) Correlation between measured ΔChIP and ΔChIP predicted by models based on motif presence changes, denoted by ‘pΔChIP(ΔSTAP)’. For each TF-TP condition, average Pearson correlation coefficient from 5-fold cross-validation is reported. (B) Similar to (A), except that the ΔChIP predictions are now based on changes in accessibility, denoted by ‘pΔChIP(ΔAcc)’. These values are similar to those reported in Figure 2B, but with slightly modified models (see text). (C) Comparison of motif-based models (x-axis) and accessibility-based models (y-axis). P-values of Pearson correlation coefficient (r) with sample size of 2754 are also shown. (D,E) Predictions of ΔChIP based on both motif changes and accessibility changes, denoted by pΔChIP(ΔSTAP+ΔAcc)’, are better than using only motif changes (D) or only accessibility changes (E).

We next sought to compare the accuracy of ΔSTAP-based predictions of ΔChIP to that of ΔAcc-based predictions, with the intention of assessing the relative contributions of sequence- and accessibility-level changes to TF binding change between species. For this, we modified the accessibility-based predictor introduced above, which used the accessibility scores for the time point matching the ChIP data set, to now use data from all three time points with available data. This allowed us to predict ΔChIP scores even for the two time points – TP2 and TP4 – for which accessibility data were not generated, by basing those predictions on accessibility scores from TP1, TP3 and TP5 (See Supplementary Figure S5A for clarification about a potential methodological concern that this might raise). Correlation coefficients between predicted and measured ΔChIP scores (Figure 3B) had an average value of 0.29 across the 14 TF:TP conditions, which is comparable to the 0.30 average correlation seen above with motif-based predictors (Figure 3A), though there is a greater variation across TF:TP conditions when using accessibility-based predictors.

We then made direct comparisons between motif-based and accessibility-based predictors of ΔChIP scores for every TF:TP condition (Figure 3C). In some cases, e.g., Twi at TP1 and Tin at TP2, changes in accessibility shows better predictive power than changes in motifs (PCC values of 0.45 vs. 0.3 for Twi:TP1 and 0.44 vs. 0.33 for Tin:TP2). This is unlikely to be due to inferior motifs used in the STAP models, as the single species STAP models for both Twi and Tin show strong correlations with ChIP (Supplementary Figure S4). It may be in part because DNA-binding of these two TFs is believed to depend not only on their own motif but also on co-binding with each other (Khoueiry, et al. 2017). In other cases, such as Bin (at all three time points), change in motif presence is a far better predictor of binding change than are changes in accessibility. This is in concordance with our previous studies in a single species – Bin motifs are very predictive of Bin binding (Junion, et al. 2012; Khoueiry, et al. 2017). For Mef2, the only TF expressed and with ChIP measurements at all five time points, ΔChIP values at later time points are predicted better using the motif-based predictor and earlier ΔChIP values are better predicted using accessibility changes, even though the motif used is the same in all cases. Interestingly, we note that this is part of a general trend for accessibility-based predictions to be better at earlier time points than later ones, such as TP4 and TP5 (Supplementary Figure S5B). This trend may be due increased embryo heterogeneity at later developmental stages having a distortive effect on cell type specific accessibility seen in bulk whole embryo DHS measurements or alternatively due to pioneering roles of early TFs priming enhancers for activation at later stages of embryogenesis.

Having found that the contribution of ΔAcc to ΔChIP is similar in extent to the contribution of ΔSTAP (change of motif presence) to ΔChIP, we asked if combining these two pieces of information would further improve our ability to predict binding changes; this would imply that divergence in accessibility and sequence have complementary information regarding change of binding. Generally, the answer was affirmative: for almost all TF:TP pairs the ΔChIP scores are better predicted with combined models (SVR using *D. melanogaster* STAP, ΔSTAP, Acc and ΔAcc features), achieving correlations in the range of 0.3 to 0.5, with an average of 0.4 across the 14 data sets, compared to ~0.3 when using accessibility or sequence alone (Figures 3D, 3E). As above, we asked how well the combined model-predicted ΔChIP values classify the enhancer pairs with the greatest increase in TF binding versus those with the greatest decrease in binding (ΔChIP in top and bottom 10 percentile respectively). We noted an AUROC of 0.84 or greater on such balanced data sets for four of the 9 TF:TP pairs (Supplementary Figure S6). The performance of this joint predictor is a quantitative summary of how well we understand the determinants of TF-binding changes between orthologous enhancers in a well-studied regulatory system.

Our results suggest that the two types of information (motif and accessibility) are complementary in their contribution to predicting changes in binding (most points are above the diagonal in Figure 3D,E). For instance, the strongest correlation observed with the joint predictor is for TWI-TP1, with a PCC of 0.52, compared to 0.3 when using motif change alone and 0.45 when using accessibility change alone. The only exceptions are data sets for Bin, where predictions of occupancy change based on sequence changes are nearly unaffected after adding accessibility information (Figure 3D), which implies that motif change alone is a strong predictor of Bin occupancy divergence. We note that in order to make such direct comparisons between determinants of binding change, we have used an approach that goes beyond testing statistical enrichments of various events, such as motif loss or gain, in regions of binding change.

### A strategy to assess predictions of binding change relevant to enhancer activity

In the analysis above, we quantified the ability to predict changes in binding by directly correlating experimentally measured ΔChIP of a TF with computationally predicted ΔChIP based on accessibility and sequence-level changes between orthologous enhancers. What does this imply for one of the ultimate goals of comparative cis-regulatory profiling – to predict changes in enhancer-driven expression? Prior work has shown that one can predict spatio-temporal activity of mesoderm enhancers based on ChIP data for the set of five TFs studied here (Zinzen, et al. 2009). We asked therefore if our ΔChIP predictions agree with the experimentally measured ΔChIP values when examined through the lens of such an activity prediction model, rather than through direct correlations for each TF separately. In other words, if we knew the ChIP values in an enhancer, and the sequence and accessibility changes between it and an orthologous enhancer, can we predict ChIP values in the ortholog *and* use them to determine if the enhancer’s spatio-temporal activity is conserved? If so, it would indicate that our understanding of binding changes is accurate enough to be of predictive value. Note that such a comparison must integrate the information from ΔChIP scores for multiple TFs, rather than compare each TF:TP separately as was done above. In this sense, we now aim to assess ΔChIP predictions in a more integrative manner.

#### Outline of approach

A major obstacle in answering this question is the lack of data on changes in enhancer activity. There is a large collection of *D. melanogaster* enhancers with annotated activities (Gallo, et al. 2011; Kvon, et al. 2014; Zinzen, et al. 2009), but only a small set of *D. virilis* enhancers whose activities were tested experimentally (in transgenic *D. melanogaster* embryos) (Khoueiry, et al. 2017). Moreover, this small set of experimentally characterized *D. virilis* enhancers mostly exhibited conserved activity (Khoueiry, et al. 2017), exacerbating the analysis of functional changes. We therefore devised a modeling-based approach to the above question, that can be briefly described as follows (Figure 4): (1) Train a model ‘A’ that predicts an enhancer’s activity from its ChIP profile (binding levels for relevant TFs), similar to prior work (Zinzen, et al. 2009). (2) Use model ‘A’ to predict the activities of orthologous *D. melanogaster* and *D. virilis* enhancers, using their respective ChIP profiles, thus characterizing the change in (predicted) activity between the orthologs. (3) Separately, use the ChIP profile of the *D. melanogaster* enhancer and motif and/or accessibility-based predictions of ΔChIP, to *predict* the ChIP profile of the *D. virilis* ortholog. Once again, estimate the activity change between orthologs, but now relying on the predicted ChIP profile of the *D. virilis* ortholog. (4) Compare the activity changes computed in steps (2) and (3), that utilize, respectively, direct ChIP measurements or predicted ChIP profiles in *D. virilis.* The extent to which these changes agree with each other will reveal how well motif and accessibility-based predictions of ΔChIP agree with real ΔChIP when seen through the lens of enhancer function.

**Figure 4.**
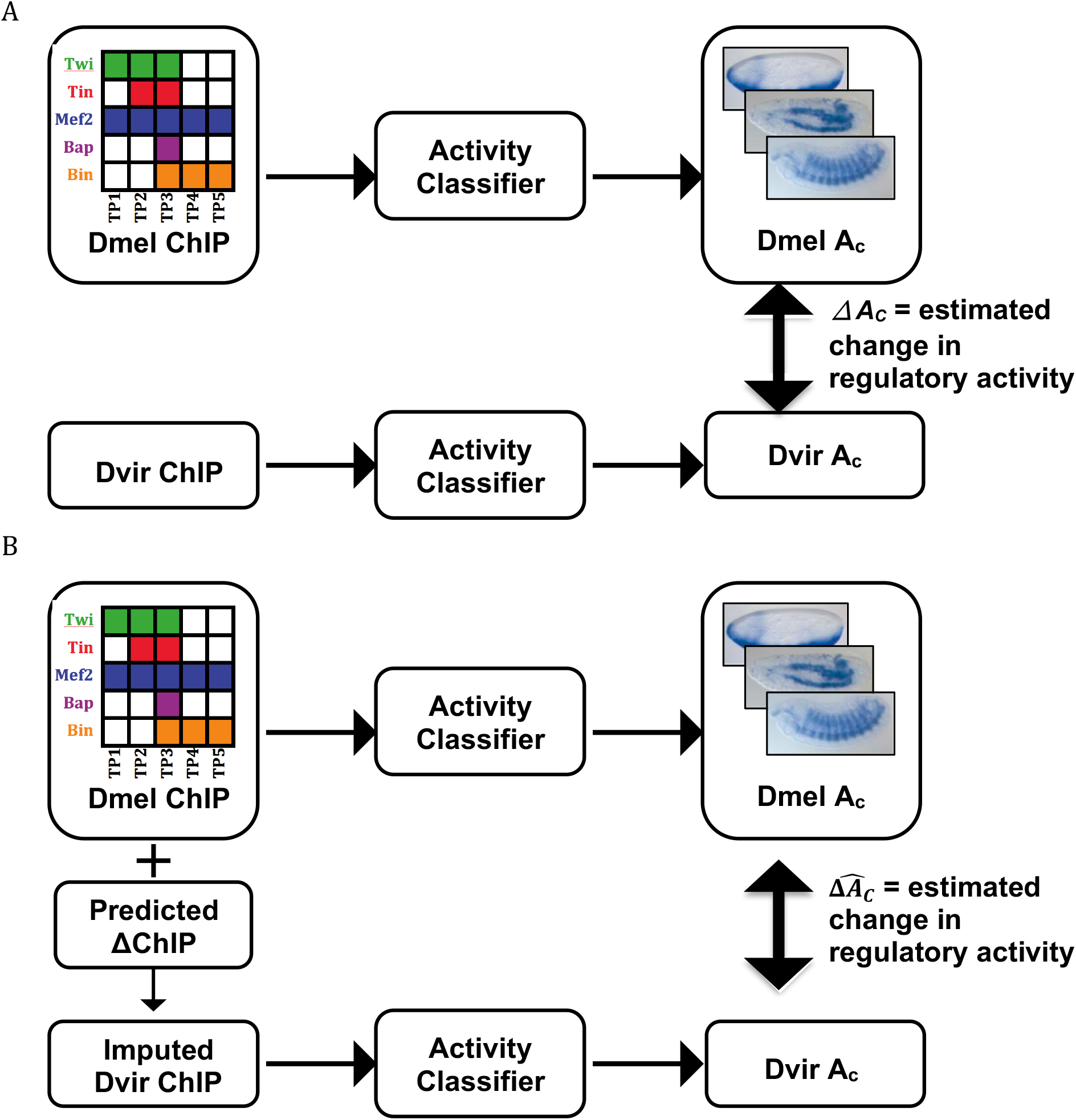
A strategy to assess predictions of binding change through the lens of enhancer activity. (A) Change in regulatory activity between orthologous enhancers is estimated from difference between output scores of activity classifiers that use *D. melanogaster* and *D. virilis* ChIP profiles respectively as input. (B) An alternative estimate of change in regulatory activity between orthologous enhancers, similar to strategy in (A), except that *D.* virilis activity classifier uses ‘imputed’ *D. virilis* ChIP profiles as input. Imputation of *D. virilis* ChIP scores is based on *D. melanogaster* ChIP scores and ΔChIP scores predicted from motif- and/or accessibility-level interspecies changes.

### Computationally imputed ChIP profiles agree with measured ChIP profiles in terms of their predictions of enhancer activity changes

We first trained XGBoost (Chen and Guestrin 2016) classifiers to predict enhancer activity in *D. melanogaster* from the 14-dimensional vector of ChIP scores of the enhancer, the ChIP scores pertaining to the 14 TF:TP conditions. Here, enhancer activity is one of the three spatio-temporal expression classes — the early unspecified mesoderm (‘Meso’), somatic muscle (‘SM’) and visceral muscle (‘VM’) – and for each class ‘C’ a separate classifier was trained to predict the enhancer’s activity in that class (‘A_C_’) on a scale of 0 to 1, representing the confidence of that classifier. More details on building and evaluating the classifiers are provided in Methods and Table 1. We then predicted the activity A_C_ of the *D. virilis* orthologs using the same classifiers and regarded the difference between these two A_C_ values (Δ*A_C_* = A_C_(D.mel) – A_C_(D. vir)) as an estimate of the change in regulatory activity, specific to class C, between the orthologous enhancers.

**Table 1.** Classifiers trained from combinatorial transcription factor binding data can accurately predict enhancer activities. Balanced accuracy from leave-one-out cross-validation is shown for models built for each activity class: mesoderm (‘Meso’), visceral muscle (‘VM’), and somatic muscle (‘SM’). Models were trained (and tested) on 223 experimentally characterized enhancers in *D. melanogaster*; for each activity class, enhancers with that activity were positives while enhancers of the other two classes were negatives. The numbers of correctly and incorrectly classified enhancers for each model are listed. TN: true negative, FN: false negative, TP: true positive, FP: false positive.

|  | **Meso** | **VM** | **SM** |
| --- | --- | --- | --- |
| **TN** | 100 | 145 | 144 |
| **FN** | 13 | 21 | 16 |
| **TP** | 89 | 44 | 50 |
| **FP** | 31 | 23 | 23 |
| **Balanced Accuracy** | 0.82 | 0.77 | 0.81 |

Next, for each spatio-temporal class ‘C’, we considered all *D. melanogaster* enhancers with predicted activity A_C_ (for that class) in the top 20 percentile, i.e., enhancers with ChIP profiles that are most suggestive of activity in class ‘C’. We further restricted ourselves to the subset of these that exhibited the highest and lowest ΔA_C_ values, i.e., enhancer pairs whose ΔChIP scores are most strongly indicative of activity change (high ΔA_C_) or conservation (low ΔA_C_). We asked how well these two subsets of orthologous enhancer pairs, with the greatest and least predicted changes in enhancer activity, can be discriminated based on predicted changes of TF binding. To this end, we obtained an ‘imputed’ ChIP profile of the *D. virilis* ortholog, by using the *D. melanogaster* ChIP scores and ΔChIP scores predicted from interspecies changes in motif presence, accessibility, or both (Figure 4B), re-estimated the regulatory activity A_C_ of the *D. virilis* ortholog based on this imputed ChIP score profile, and computed its difference from the regulatory activity of the *D. melanogaster* enhancer (Inline) as an alternative estimate of change in regulatory activity between the orthologs. Finally, we computed the Pearson correlation coefficient between the two estimates ΔA_C_ and (Inline), across all enhancer pairs considered (Table 2A), and also noted that AUROC values when (Inline) is used to classify enhancer pairs with high ΔA_C_ versus low ΔA_C_ (Table 2B and Supplementary Figure S7).

**Table 2.** Changes in motif presence and accessibility can be used to predict enhancer activity change. (A) Pearson correlation coefficients between two different estimates of activity change: ΔA_C_, based on measured *D. virilis* ChIP score profiles and (Inline), based on *D. virilis* ChIP score profiles imputed from *D. melanogaster* scores and predictions of binding change (ΔChIP), which in turn were made from changes in sequence (‘ΔSTAP), accessibility (‘ΔAcc’) or both. As a random control baseline, we used *D. virilis* ChIP scores imputed from from *D. melanogaster* scores and a permuted version of the ΔChIP matrix. (B) AUROC values representing how well (Inline) values can classify high versus low ΔA_C_ enhancer pairs. These analysis were performed for three expression domains: mesoderm (Meso), visceral muscle (VM), and somatic muscle (SM). Enhancer pairs that exhibited the highest and lowest ΔA_C_ values (in top and bottom 10 percentile for classes ‘Meso’ and ‘VM’, and in the top and bottom 5 percentile for class ‘SM’) are reported.

| **D. virilis ChIP imputation**<br>**based on:** | **Meso** | **VM** | **SM** |
| --- | --- | --- | --- |
| $\Delta\text{STAP}$ | 0.33 | 0.42 | 0.36 |
| $\Delta\text{Acc}$ | 0.21 | 0.30 | 0.33 |
| $\Delta\text{STAP}$ and $\Delta\text{Acc}$ | 0.30 | 0.47 | 0.37 |
| Random control | 0.07 | 0.16 | 0.09 |
| #Samples | 110 | 110 | 56 |

(B)
| **D. virilis ChIP imputation**<br>**based on:** | **Meso** | **VM** | **SM** |
| --- | --- | --- | --- |
| $\Delta\text{STAP}$ | 0.65 | 0.70 | 0.82 |
| $\Delta\text{Acc}$ | 0.57 | 0.68 | 0.70 |
| $\Delta\text{STAP}$ and $\Delta\text{Acc}$ | 0.65 | 0.75 | 0.78 |
| Random control | 0.56 | 0.59 | 0.57 |
| #Samples | 110 | 110 | 56 |

We noted that when *D. virilis* ChIP score profiles are imputed based on motif and accessibility changes together, the two estimates of activity change have a correlation of 0.47 (P-value 2.21E-7) for the ‘VM’ class, which is substantially greater than the correlation of 0.16 (P-value 0.09) seen in a random control. (In the control setting, *D. virilis* ChIP scores were imputed based on a random permutation of ΔChIP scores). The high level of agreement between ΔA_C_ and (Inline) is also reflected in the classification AUROC of 0.75, compared to the random control AUROC of 0.59. Similarly, for the ‘SM’ class, a high agreement between the two estimates is borne out by an AUROC of 0.78 (compared to 0.57 in random control), and a correlation coefficient of 0.37 (P-value 0.005), while the random control yields a correlation of 0.09 (P-value 0.51) for this expression class. The correlation and AUROC values are lower for the class ‘Meso’, although clearly statistically significant, e.g., correlation of 0.33 (P-value 0.0004) compared to random correlation of 0.07 (P-value 0.47). Taken together, these results suggest that the accuracy of ΔChIP predictions demonstrated above (Figure 3D,E), based on modeling interspecies changes in sequence and accessibility, is sufficient for us to make similar predictions of enhancer activity changes as can be made using experimental knowledge of binding changes. It also indicates that much of variation in TF occupancy not predicted by accessibility or sequence may not be critical for fitness related biological output. At the same time, this ability to predict activity changes differs from one expression class to another and there is substantial room for improvement.

We also repeated the above analysis using imputed ChIP score profiles in *D. virilis* from ΔChIP predictions based only on sequence-level changes or only on accessibility changes, rather than both. Our main observation is that sequence-based predictions of binding change are often close to and in some cases even better than the joint predictors that utilize sequence and accessibility changes (Table 2, rows ‘ΔSTAP’ compared to rows ‘ΔSTAP and ΔAcc’). A noteworthy data point is that for the ‘SM’ class, sequence-based predictions of ΔChIP can accurately predict, with an AUROC of 0.82, the enhancer pairs with greatest and least activity change, where activity is defined based on real ChIP profiles in the two species. We also noted that ΔChIP predictions based on accessibility changes alone are consistently worse in terms of the resulting agreement between *ΔA_C_* and (Inline). The is in contrast to the observations in Figure 3C, where we did not observe a consistent difference between sequence-based and accessibility-based predictors of binding change for individual TF:TP pairs. This is not surprising: accessibility changes are indeed an important statistical determinant of binding changes, but predicting activity change likely requires correctly predicting binding changes of multiple TFs, and the sequence-based predictors have an advantage in this respect as they use different motifs for each TF, while the accessibility-based predictors utilize the same underlying information in predicting binding change for every TF.

## DISCUSSION

We examined the evolution of DNA accessibility in two distant species, and found it to be an important determinant or correlate of inter-species changes in TF binding. It is possible that changes in accessibility are not causal of binding change but rather a consequence; for instance, the relaxation of selection pressure resulting from a functional loss of TF binding may in turn lead to reduction in local accessibility, which may be the case for Twist. Interestingly, we noted that our ability to predict TF binding changes simply based on accessibility changes rivals our ability to make those predictions based on sequence divergence, i.e., change of TF motif presence. At the same time, there is substantial complementarity between the two, and a model that combines both motif and accessibility changes can predict changes in TF binding more accurately than either alone. A noteworthy feature of our work is that we have approached issues of *cis*-regulatory divergence in a quantitative manner, asking ‘to what extent’ a relationship (e.g., between accessibility and binding changes) is supported by data, in addition to asking if ‘there exists strong evidence’ for such a relationship, through hypothesis testing approaches. Such a quantitative approach is also important for comparing how well two different types of information – changes in accessibility and motif presence – correlate with binding change.

While most of our analyses were aimed at global insights, we also made several TF-specific observations that suggest properties of those TFs. For instance, the relatively high concordance between accessibility change and binding change for Twi suggests to us that Twi may have direct influence in making DNA accessible (also suggested in (Cheng, et al. 2013; Sandmann, et al. 2007)), though other explanations are also feasible. Another analysis revealed that the co-divergence of Twi and Tin binding sites (Khoueiry, et al. 2017) may only partly be explained by the shared influence of accessibility, thus increasing the support for alternative causes such as extensive cooperative binding, evidence for which was also reported in (Cheng, et al. 2013). This latter possibility was also consistent with our observation that motif-level changes were less effective than accessibility changes in predicting binding divergence of either TF.

It is worth emphasizing that our comparisons of sequence, accessibility and TF binding between species have the advantage of being performed in the context of a system where the examined TFs are all essential regulators that participate in a highly interconnected regulatory network, participating in feed-back and feed-forward regulation of a large number of genes. Thus, by focusing on putative enhancers defined by multiple ChIP peaks in close proximity, we hope to have enriched for evolutionary events with potential consequences for gene expression. Such an advantage is often not possible in other studies of binding evolution, since few regulatory networks have been as well characterized (see (Paris, et al. 2013) for another example).

We also examined how total changes in TF binding relate to changes in enhancer activity within the mesoderm specification network. We found most orthologous enhancers have conserved activity despite high divergence in TF binding events (see details in Supplementary Text S1). Enhancers in this network have been previously shown (Zinzen, et al. 2009) to be amenable to computational models that predict their activity (tissue specificity) from their TF binding profiles within one species (*D. melanogaster*). It was thus natural to ask if evolutionary changes in TF binding can be interpreted in the light of such functional models. However, we were unable to answer this question in the most direct way– whether binding changes for multiple TFs can, via these models, predict changes in enhancer activity – because the available data on regulatory activities of orthologous enhancers are sparse. Instead, we used the ability to model enhancer activity from ChIP data to show that predicted changes in binding (based on accessibility and motif divergence) agree with measured binding changes (ChIP data) in terms of what they imply about activity changes. It is worth clarifying that we defined activity change between orthologous enhancers as the difference in predicted activity in a spatio-temporal class, using a computational model that is meant to predict enhancer activities in *D. melanogaster*. Thus, under this definition, the activity of a *D. virilis* enhancer is in fact the expression pattern we predict it to drive if it was tested through a reporter assay in a *D. melanogaster* embryo. This was necessary since we do not yet have sufficient training data (*D. virilis* enhancers with known expression readouts in *D. virilis*) to learn a classifier for predicting activity in *D. virilis*. It was also a convenient choice since we did not have to make assumptions about conservation of the *trans* context between *D. melanogaster* and *D. virilis*. The only assumption required was that a “ChIP profile” (14 TF-ChIP values at a enhancer) obtained from *D. virilis* is semantically comparable to a ChIP profile obtained from *D. melanogaster*, which we previously showed is the case (Khoueiry, et al. 2017). Also, the absolute values in the ChIP profile do not matter (only their relative values), since we worked with normalized ChIP profiles, which have similar distributions in both species (Supplementary Figure S8). There is precedence in the literature for examining activity changes between orthologous enhancers in a common cellular context (Arnold, et al. 2014). Expression is often conserved despite divergence at sequence level, but that the data is sparse still. We expect that more experimental data on heterologous activity, e.g., of *D. virilis* enhancers, will better address the functional consequences of binding changes and improve our ability to predict functional cis-regulatory change from accessibility and sequence data.

In ending, we note that even when using a combined model that integrates sequence and accessibility data, we were able to predict TF binding change with a correlation coefficient of ~0.5 at best. What is missing in the data and models that might account for the missing predictability? The answer is probably closely tied to the same issue in the context of single-species TF binding prediction, a topic that has received far greater attention (Slattery, et al. 2014), and where a number of additional factors, such as co-binding and competitive binding (Cheng, et al. 2013; Wasson and Hartemink 2009), more precise motif characterizations (He, et al. 2009; Weirauch, et al. 2013), higher resolution mapping of chromatin context (Cheng, et al. 2013; Peng, et al. 2015; Pique-Regi, et al. 2011), etc. have been shown to improve predictive ability. Incorporation of these additional dimensions of data and modeling in the future should further increase our understanding of evolutionary changes in transcription factor binding.

## MATERIALS AND METHODS

### ChIP data and enhancers

We collected TF-ChIP data on five developmental TFs across five stages of embryogenesis, in the form of ChIP-chip in *D. melanogaster* (Zinzen, et al. 2009) and ChIP-seq in *D. virilis* (Khoueiry, et al. 2017), from previous studies. The five TFs examined were Twist (Twi), Myocyte enhancer factor-2 (Mef2), Tinman (Tin), Bagpipe (Bap) and Biniou (Bin) (Figure 1A). A total of 14 TF-time point pairs, ‘TF:TP conditions’ or simply ‘conditions’, were analyzed in this study (Figure 1B). ChIP peaks in close proximity across all TF:TP conditions were clustered to define 8,008 putative ChIP enhancers in *D. melanogaster* (Zinzen, et al. 2009) and 10,532 putative ChIP enhancers in *D. virilis* (Khoueiry, et al. 2017). A ChIP score was then assigned to each enhancer, for each TF:TP condition, by extracting the mean ChIP signal (using library size normalized bigwig files) over the enhancer boundaries (performed in Galaxy (Afgan, et al. 2016) using the “Compute mean/min/max of intervals” tool version 1.0.0). To make ChIP scores comparable across stages and species, we applied the following normalization on ChIP scores for each TF:TP condition: we set μ+3σ as the maximum ChIP score, where μ and σ are the mean and standard deviation across all putative enhancers, replaced all ChIP scores greater than this maximum with μ+3σ, and finally applied min-max normalization to set all ChIP scores in a range between 0 to 1.

Orthologous enhancer pairs were defined in our previous study (Khoueiry, et al. 2017). Briefly speaking, we translated *D. virilis* enhancer coordinates into *D. melanogaster* coordinates, overlapped the 10,532 *D. virilis* enhancers with the 8,008 *D. melanogaster* enhancers, and finally obtained a set of 2754 orthologous enhancer pairs. This set of orthologous enhancers served as the subjects of analysis in this study.

### DNase-seq sample processing

Accessibility data in *D. virilis* and *D. melanogaster* were obtained using DNase-seq from whole embryos at developmental stages 5-7, 10-11, and 13-15, referred to as TP1, TP3, and TP5. The developmental stages of timed collections were determined exactly as described in (Khoueiry, et al. 2017). Raw paired-end reads were aligned using BWA (Li and Durbin 2009) on Flybase-R1.2 assembly version for *D. virilis* and on Flybase Assembly 5 (dm3) for *D. melanogaster*. Reads were filtered for optical and PCR replicates using samtools (Li, et al. 2009). For peak calling, we used MACS2 (–to-large with -g 1.2E8 for *D. melanogaster* and 1.9E-8 for *D. virilis* and -p 1E-3 as requested for the Irreproducibility Discovery Rate analysis, or IDR (Landt, et al. 2012)). We derived peaks using 1% IDR threshold leading to a unique highly confident and consistent peak sets for biological replicates. For visualization and generation of bigwig score files, reads from BAM files were extended to the average length of the genomic fragments for the corresponding time point, merged and scaled to Read Per Million (RPM) using deeptools (Ramírez, et al. 2014). Each enhancer (in either species) was assigned an ‘accessibility score’ by extracting the mean DNase signal (abovementioned bigwig files) over the enhancer boundaries (performed in Galaxy (Afgan, et al. 2016) using the “Compute mean/min/max of intervals” tool version 1.0.0). To make accessibility scores comparable across stages, for each time point, we performed the same normalization as we did for ChIP scores, i.e. replacing accessibility scores greater than μ+3σ with μ+3σ, where μ and σ are the mean and standard deviation, and then applying min-max normalization.

### Support Vector Regression models to predict changes of ChIP scores

For every TF:TP condition, we trained Support Vector Regression (SVR) models, using the R package ‘e1071’ (Meyer, et al. 2015), to predict the interspecies differences in ChIP scores, defined as ΔChIP=ChIP_Dmel_-ChIP_Dvir_ for each enhancer. We used the set of 2,754 orthologous putative enhancer pairs to train and evaluate models. For each orthologous pair, two kinds of input features were used to predict ΔChIP: 1) *D. melanogaster* accessibility score and interspecies accessibility score changes (ΔAcc=Acc_Dmel_-Acc_Dvir_) for the appropriate time point, and 2) *D. melanogaster* STAP score and interspecies change in STAP scores (ΔSTAP=STAP_Dmel_- STAP_Dvir_) for the appropriatew TF:TP condition. STAP scores represent motif-based prediction of TF occupancy in a segment of sequence, and have a free parameter representing TF concentration, trained on ChIP data for the TF:TP condition (see subsection below for details). Separate models were trained, that used (a) only accessibility features (‘accessibility-based model’), (b) only motif features (‘motif-based model’), or (c) both types of features (‘combined model’). To have a direct comparisons between motif-based and accessibility-based predictors of ΔChIP scores for every TF:TP condition (Figure 3), we modified the accessibility-based SVR models by using data from all three time points simultaneously. Similarly, features of the combined SVR model included *D. melanogaster* STAP and ΔSTAP at the matching TF:TP condition, as well as *D. melanogaster* Acc and ΔAcc for all three time points. SVR models were trained with default parameters. Performance was measured by Pearson correlation coefficient between measured and model-predicted ΔChIP scores, using 5-fold cross-validation.

### STAP models to predict TF occupancy based on motif presence

STAP (Sequence To Affinity Prediction) program (He, et al. 2009) is a thermodynamics-based model that integrates one or more strong as well as weak binding sites, using a given motif, to predict net TF occupancy within a DNA segment. For each TF:TP condition in each species, we trained a STAP model, following the procedures we used previously in Cheng et al. (2013). We chose the top 1,000 ChIP peaks as the positive training set and 1,000 random windows of the same length as the negative training set, along with their respective normalized ChIP scores. ChIP peaks overlapped with the orthologous enhancers were excluded in training set. The binding motif (position weight matrix, PWM) for each TF was based on the best performing PWMs discovered from *D. melanogaster* and *D. virilis* ChIP data (Khoueiry, et al. 2017) (Supplementary Figure 1). A single free parameter of STAP was learned based on this training set.

To assess the performance of STAP model on each of the 28 ChIP data sets in the given TF, time point, and species combination, we applied four-fold cross-validation on the 2,000 DNA segments training set. Each fold used 1,500 DNA segments to train the single free parameter in STAP, and 500 DNA segments to score. The resulting 2,000 STAP scores, aggregated from each fold, were compared to respective ChIP scores, by Pearson correlation coefficient (PCC). These 28 STAP models, previously reported in (Khoueiry, et al. 2017), fit the ChIP data well, and showed an average PCC of 0.51. We also checked the single parameter of STAP learned in each fold, and observed similar values across four folds.

Once the STAP model was trained for every TF, time point, species combination, we used the STAP model to score each enhancer for motif presence. STAP scores were further normalized in the same way as ChIP scores, i.e. capping outliers at μ+3σ, where μ and σ are the mean and standard deviation, and then applying min-max normalization.

### Experimentally characterized enhancers

To build a training set for the enhancer activity classifier, we collected known mesoderm enhancers from our previously built CRM Activity Database (CAD) (Zinzen, et al. 2009), activity information of active tiles from Kvon et al. (2014), and a set of new entries from RedFly database (Gallo, et al. 2011). Three activity classes were considered: mesoderm (Meso), visceral musculature (VM), and somatic musculature (SM). Enhancers that drive expression in more than one classes (e.g. Meso and SM or VM and SM) were excluded. We then overlapped the annotated enhancers with our 2,754 orthologous putative ChIP enhancers in *D. melanogaster*. This led to a final training set of 233 enhancers, with 102 expressed in Meso, 65 in VM, and 66 in SM.

### XGBoost models to predict enhancer activities

XGBoost (Chen and Guestrin 2016) is a supervised machine learning method that uses training data with multiple features to predict a target variable. For each activity class ‘C’, an XGBoost classifier Ac was trained by using the R package ‘xgboost’ (Chen, et al. 2016) to discriminate between members and non-member of the class. Thus, for the Meso class, the positive set includes enhancers with Meso annotation, while the negative set includes enhancers with VM or SM annotations. The input features for each enhancer were a 14-dimensional vector of ChIP scores of that enhancer pertaining to the 14 TF:TP conditions. To adjust for the imbalanced distribution of training data set, we used the Synthetic Minority Over-sampling Technique (SMOTE) (Chawla, et al. 2002), from R package ‘DMwR’ (Torgo 2016), to oversample the minority class. We trained the XGBoost classifiers in the mode of ‘logistic regression for binary classification (binary:logistic)’. Parameters were set as below: ‘eta’ = 0.2, ‘nrounds’ = 50, ‘max_depth’ = 4, ‘subsample’ = 0.9, ‘colsample_bytree’ = 0.8, by following the guidelines from XGBoost documentation. Once we trained the activity classifier Ac, leave-one-out cross-validation was applied to measure the performance.

We trained A_C_ on 223 experimentally characterized enhancers (Gallo, et al. 2011; Kvon, et al. 2014; Zinzen, et al. 2009) associated with the three expression classes, and noted balanced accuracy values around 0.8 in leave-one-out cross validation for each class (Table 1). When estimating accuracy for any class, enhancers of that class were treated as positives, and enhancers of the other two classes were considered as negatives. For each classifier, the specificity is ~0.9 and sensitivity is ~0.7. We also assessed the accuracy of the trained functions on held-out transgenic reporter assays of *D. melanogaster* and *D. virilis* enhancers (Khoueiry, et al. 2017; Zinzen, et al. 2009). Among 35 experimentally tested enhancers, the predictions of 23 were correct (drove expression in the predicted domain), 3 were partially correct (one of the active tissues was predicted), whereas 9 predictions failed (did not drive any expression in the predicted domain) (Supplementary Table S2). The experimental assays comprised of enhancers in both species, and the accuracy noted in these evaluations justified our assumption that classifiers trained in *D. melanogaster* can be used to predict the activities of *D. virilis* enhancers as well (though in a *D. melanogaster* context).

We noted that similar enhancer activity predictors had been presented in Zinzen et al. (2009), where Support Vector Machines (SVMs) trained from ChIP scores were shown to accurately predict enhancer activities in *D. melanogaster*. We rebuilt the classifiers here mainly because our desired tradeoff between sensitivity and specificity was different; in particular, we sought to achieve high values of balanced accuracy when evaluating classifiers on imbalanced data sets (in our case, there are more negative samples than positive samples); see Supplementary Table S3. In addition, ChIP data for Tin at TP1, an input feature for *D. melanogaster* activity classifiers reported in (Zinzen, et al. 2009), was not available for *D. virilis* (Khoueiry, et al. 2017), further necessitating rebuilding of classifiers.

### NGS data availability

Raw sequence data has been deposited in ArrayExpress under accession numbers E-MTAB-3797 (*D. melanogaster* and *D. virilis* DNAse developmental time courses)

